# Predicting gene expression from cell morphology in human induced pluripotent stem cells

**DOI:** 10.1101/2022.04.19.488786

**Authors:** Takashi Wakui, Mitsuru Negishi, Yuta Murakami, Shunsuke Tominaga, Yasushi Shiraishi, Anne E. Carpenter, Shantanu Singh, Hideo Segawa

## Abstract

Purification is essential before differentiating human induced pluripotent stem cells (hiPSCs) into cells that fully express particular differentiation marker genes. High-quality iPSC clones are typically purified through gene expression profiling or visual inspection of the cell morphology; however, the relationship between the two methods remains unclear. We investigated the relationship between gene expression levels and morphology by analyzing live-cell phase-contrast images and mRNA profiles collected during the purification process. We employed this data and an unsupervised image feature extraction method to build a model that predicts gene expression levels from morphology. As a benchmark, we confirmed that the method can predict the gene expression levels from tissue images for cancer genes, performing as well as state-of-the-art methods. We then applied the method to iPSCs and identified two genes that are well-predicted from cell morphology. Although strong batch effects resulting from the reprogramming process preclude the ability to use the same model to predict across batches, prediction within a reprogramming batch is sufficiently robust to provide a practical approach for estimating expression levels of a few genes and monitoring the purification process.

## 1. Introduction

Human induced pluripotent stem cells (hiPSCs) are of growing importance in both basic and translational biomedical research due to their capacity to differentiate into any cell type and proliferate indefinitely. HiPSCs are derived from easily accessible somatic cells, like leukocytes or skin fibroblasts, using different iPSC reprogramming methods ^(1,2^. Unfortunately, currently available iPSC reprogramming methods are stochastic ^(3,4^, which leaves a subset of cells partially reprogrammed and with low pluripotency. Therefore, generating high-quality iPSCs requires an extensive and time-consuming purification process to eliminate partially reprogrammed cells ^(5^.

During the purification process iPSC clones with high pluripotency and stemness are selected based on either gene expression profiling data, such as RNA-seq, or cell morphology as evaluated by visual inspection or image analysis ^(6–12^. However, both of these approaches have major limitations. For example, although gene expression profiling gives a direct readout of stemness and differentiation markers ^(13,14^, the method is destructive, precluding further analysis of the cell. Therefore, the outcome can only be estimated through statistical interpretation using multiple samples, which are costly to grow and propagate. On the other hand, morphological analysis of cell images is non-destructive; however, this method identifies iPSCs with high pluripotency based on their more compact shape with less cytoplasm ^(15,16^, thus using indirect readouts prone to errors and misinterpretation, with no clear link to underlying gene expression. Therefore, an improved strategy for identifying high-quality iPSC populations that overcomes limitations of the current methodologies by bridging gene expression with morphology in a non-destructive manner would help address this major bottleneck in iPSC generation process.

Very recently, several studies have demonstrated that morphological features can be used to predict gene expression in various contexts ^(17–21^. For example, supervised deep learning models, such as a Convolutional Neural Network (CNN), were trained with datasets of pathological images and spatial gene expression profiles obtained by single-cell sequencing of the exact same sample to predict cancer gene expression from pathological images. The resulting models were able to accurately link morphology with gene expression, suggesting that a similar strategy could be extended beyond cancer and applied to the iPSC purification process. However, these strategies require expensive single-cell sequencing measurements, which limits their wide-spread adoption.

In this study, to overcome existing limitations and provide the community with a more robust, affordable, easy to implement, and less time-consuming iPSC purification strategy, we developed a method to predict the expression of iPSC genes from brightfield cell morphology during the iPSC purification process. Our method was designed to use multiple label-free, phase-contrast images of cells growing in culture vessels and bulk-level gene expression profiles of these cells, measured for each vessel. Thus, unlike most previous studies, the gene expression profiles (by virtue of being at the bulk level) do not have a one-to-one correspondence with individual cells. Therefore, we employed an approach based on an unsupervised deep learning model to extract morphology features from each image, and then created a simple Support Vector Regression (SVR) model to predict gene expression based on the image features of each vessel. We validate our approach on a benchmark dataset, as well as data obtained from iPSC samples obtained from experimental conditions designed to simulate iPSC purification process.

## 2. Results

### 2.1. Gene expression data set assembly for predictive model building

In order to build a robust predictive model, we focused our initial efforts on acquiring high-quality imaging and bulk gene expression data under conditions that simulate iPSC purification process. We generated six datasets composed of 29 iPSCs clones derived from peripheral blood mononuclear cells (PBMCs) of 15 donors. Datasets A1 to A3 were obtained by culturing 15 iPSC clones derived from a single donor, and datasets B1 to B3 were generated by culturing 14 iPSC clones derived from 14 different donors (Supplemental Table 1). To simulate different stages of iPSC purification processes, all iPSCs were cultured for three passages with increasingly-sized vessels (6-well plate, T75 flask, and T150 flasks) for each passage respectively (Supplemental Figure 1). In every passage, the cells were passaged on Day 4, and phase-contrast imaging and gene expression measurements were performed before passaging.

**Supplemental Fig. 1.**
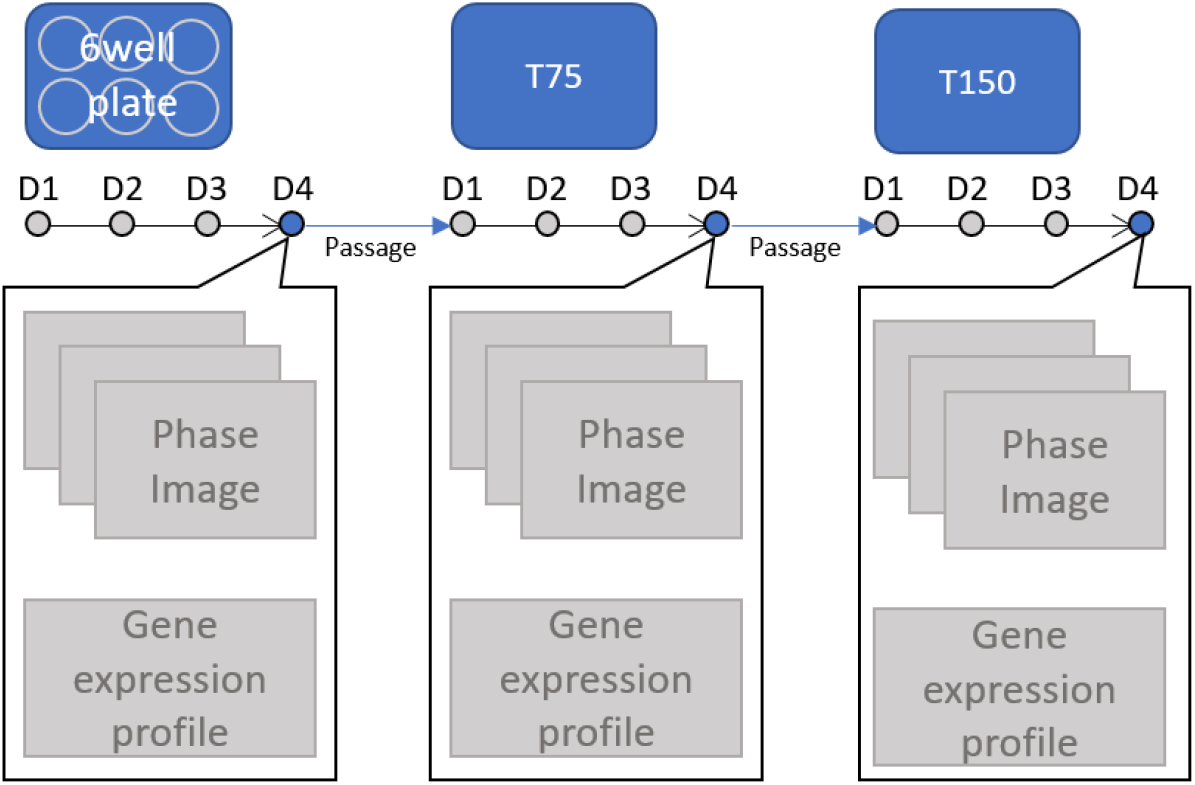
Overview of culture process. All cells were cultured for four days after passage. Phase-contrast imaging and bulk RNA-seq analysis were performed before passaging on Day 4 for each of the three stages.

**Supplemental Table 1.** Dataset structure. Datasets of experiment A (A1-A3) were obtained from the samples of 15 clones derived from one donor, while those of experiment B (B1-B3) were obtained from the samples of 14 clones derived from 14 donors.

| Experiment | Clones | Timepoint 1<br>(6 well plate,<br>D4) | Timepoint 2<br>(T75, D4) | Timepoint 3<br>(T150, D4) |
| --- | --- | --- | --- | --- |
| **A**<br>**(1 donor)** | 15 clones<br>(Clone #1-15) | Dataset A1 | Dataset A2 | Dataset A3 |
| **B**<br>**(14 donors)** | 14 clones<br>(Clone #16-29) | Dataset B1 | Dataset B2 | Dataset B3 |

Our gene expression analysis was done using Targeted Amplicon Sequencing, a bulk RNA-seq strategy focused on a subset of genes. We measured expression levels for 980 genes related to stemness and pluripotency. Out of those 980 genes, we filtered the genes by their average (mean) and coefficient of variation values (CV) across all the samples to identify genes with sufficient expression levels and significant variance, as those suggest predictive power needed for robust model building. The final list contained 218 genes, including three Yamanaka factor genes: KLF4, SOX2, and POU5F1 (Figure 1). Thus, this data represents the input gene expression data for our predictive model training.

**Figure 1.**
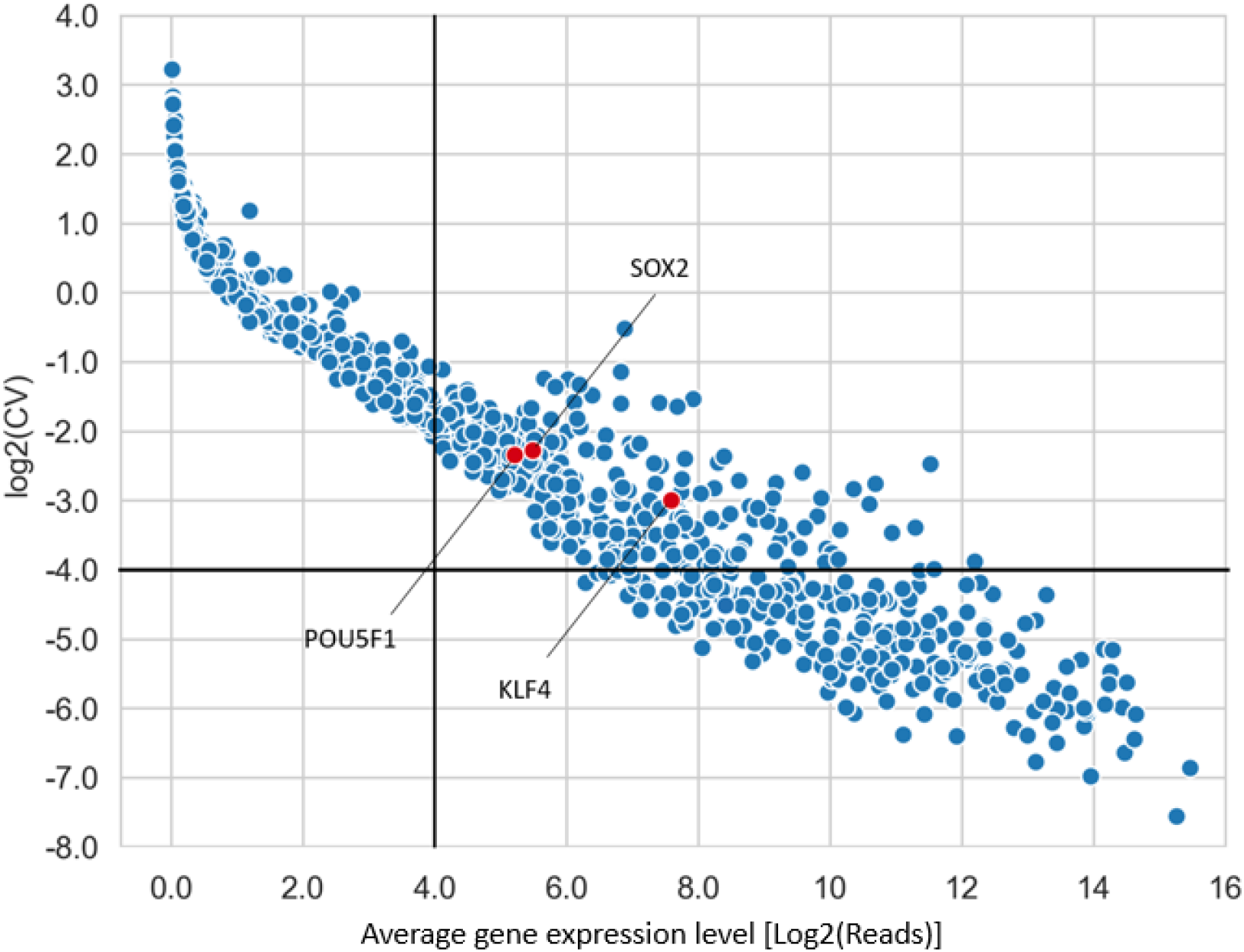
Distribution of gene expression levels and variation across samples. Each dot represents a gene expression of a single gene. After filtering based on the average (X-axis) and coefficient-of-variation (Y-axis) of the gene expression level, we retained 218 genes out of the 980 genes (top right quadrant), including three genes of particular interest (red dots).

### 2.2. Image-based gene expression prediction pipeline

To develop a robust image-based gene expression prediction pipeline that would require a limited amount of bulk level mRNA data (above), we adopted a 2-stage approach (Figure 2). First, given the number of samples (87; 29 clones at 3 different time points), we treated each individual iPSC phase-contrast image as a collection of thousands of pieces of imaging information by splitting it into patches of 160×160 px (72 μm), resulting in 1,024 patches per image. To extract features from these patches, we used a modified VQ-VAE-2 model ^(22^ as a self-supervised feature extraction method (Supplemental Figure 3). Thus, in stage 1, we extracted vector-quantized feature maps by applying the trained VQ-VAE-2 model to all the patches (see Methods for details). Next, in stage 2, we used this feature extraction process together with SVR to obtain 128-dimensional feature vectors for each image patch, which are then averaged to produce a single 128-dimensional profile for each sample.

**Figure 2.**
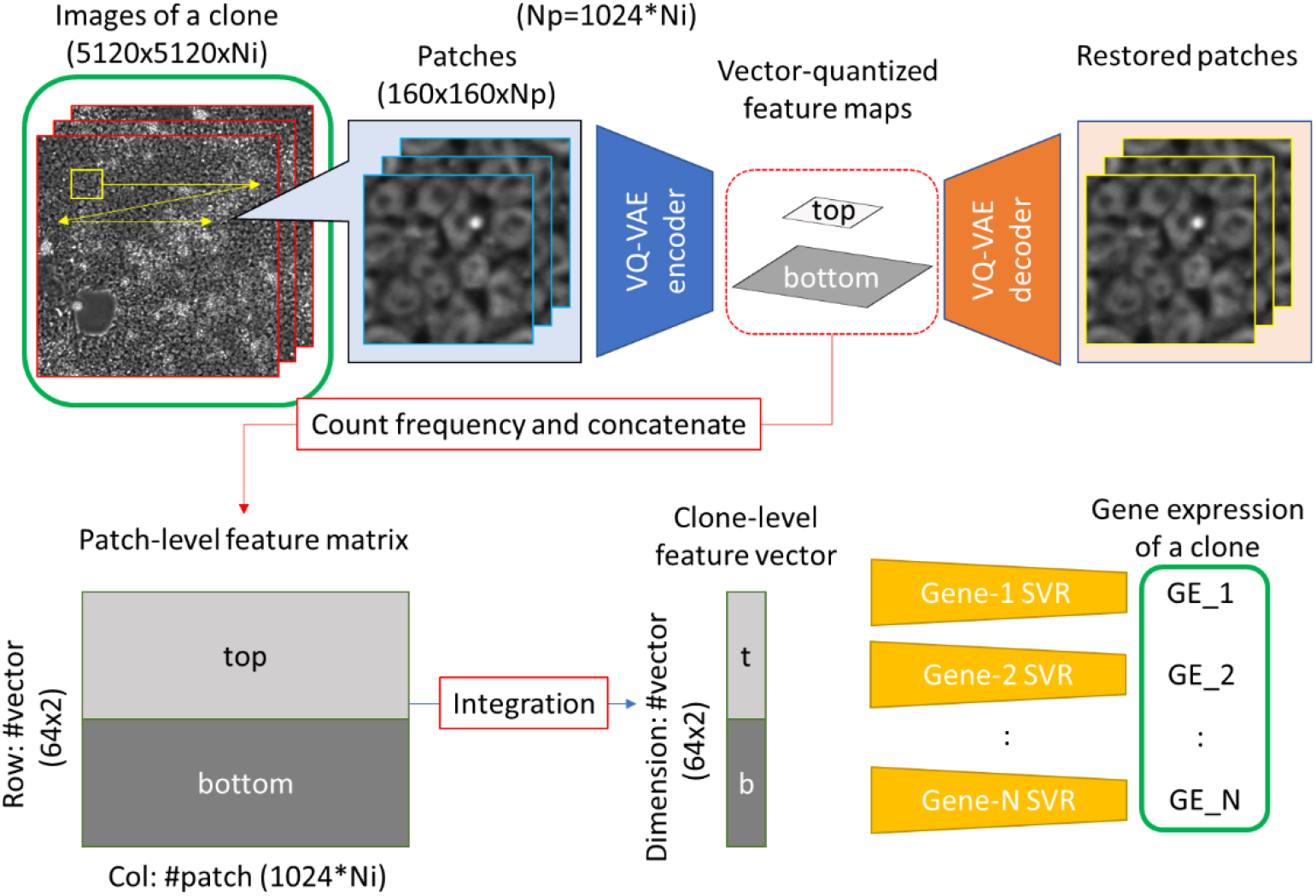
Gene expression prediction pipeline. We split iPSC phase-contrast images of a sample into patches of 160×160px (72 μm) each, extracted feature maps using the VQ-VAE encoder, and integrated the maps into a feature vector. We constructed 218 SVR models, corresponding to the 218 target genes.

**Supplemental Fig. 3.**
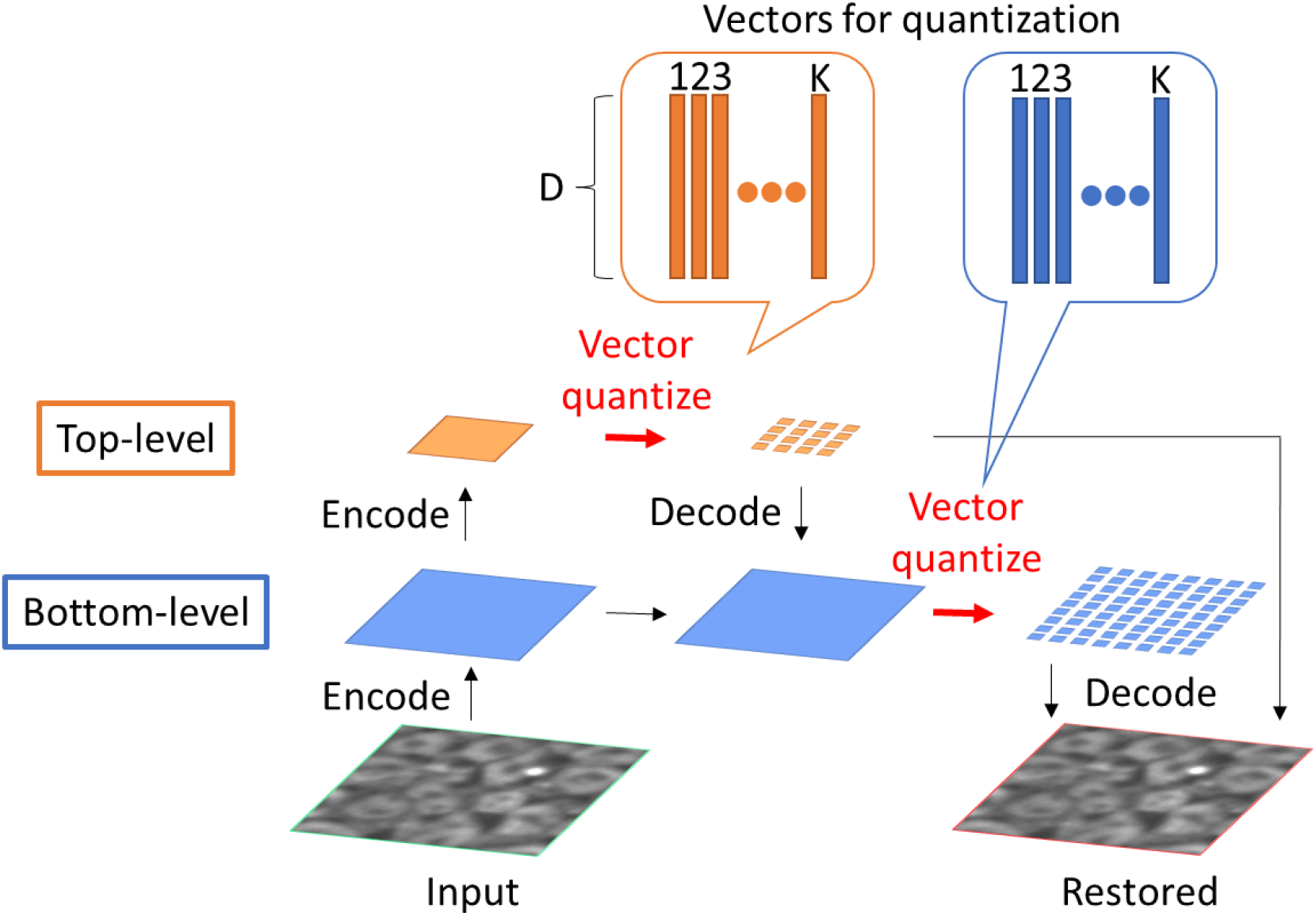
VQ-VAE-2 encoder and decoder. We encoded an input patch into two feature maps with distinct resolutions (top- and bottom-level). Each of the feature maps with two different resolutions (top- and bottom-level) encoded by CNN is quantized element-wise by the nearest one of K distinct embedding vectors of D dimension, so the vector-quantized feature maps have integers from 1 to K, which represent indices of the embedding vectors. We decoded vector-quantized feature maps into a restored patch. The vectors for quantization and the parameters of the encoder and the decoder are all trained to minimize mean square error (MSE) between input and restored patch.

### 2.3. VQ-VAE-2 model validation

To confirm that the trained VQ-VAE-2 model extracted sufficient image features to reconstruct images, we evaluated the reconstruction accuracy for each dataset using mean-squared error (MSE) between the original images and the reconstructed images (Figure 3). We used dataset A3 for the VQ-VAE-2 model training. Then, we randomly sampled 10 images (10240 patches) of each dataset to calculate MSE. All of the MSEs are lower than 36, which means that the average pixel-wise error for the 8-bit images (256 levels) is less than 6 (2.3%) for all images; little difference in the images was found by visual inspection across the datasets (Figure 3a). We thus concluded that our VQ-VAE-2 model was sufficiently trained, as it can extract the image features essential to reliably reconstruct the iPSC images.

**Figure 3.**
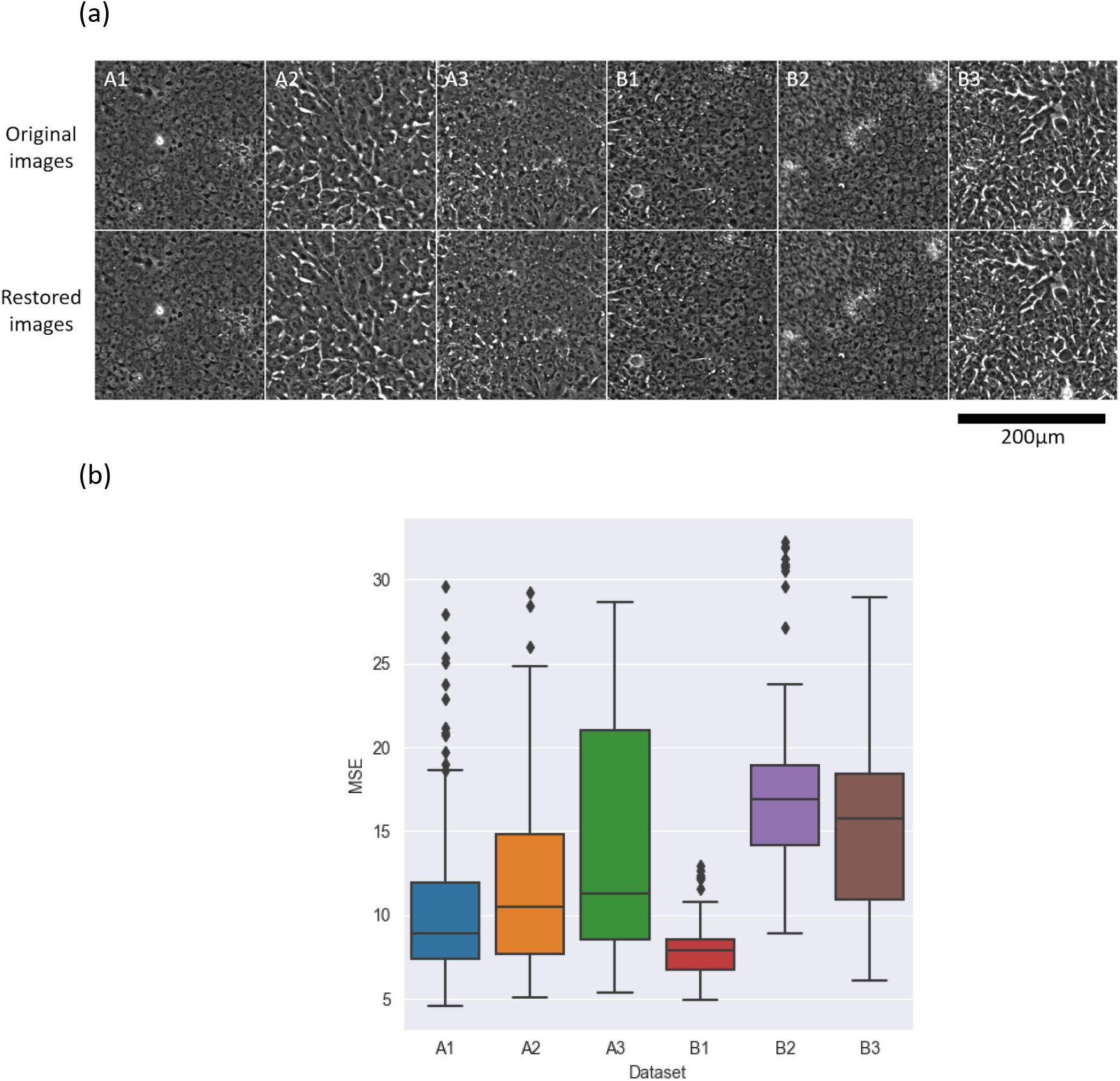
Image restoration accuracy of each dataset. The VQ-VAE-2 model was trained on a single dataset (A3) and applied to all datasets to extract image features. (a) Comparison of the original images and the restored images by the model trained with dataset A3. (b) Average MSEs calculated between the original images and the restored images (lower MSE is better). All MSEs were less than 36 (i.e. averages were less than 6) for 8-bit images, indicating good restoration performance regardless of the dataset.

### 2.4. Extracted image features relate to experimentally-defined quality categories

To examine the quality of extracted image features, we investigated the relationship between the image features and the four morphological categories (*undifferentiation, cracked, built-up,* and *differentiation)* that are routinely used by iPSC culture experts to identify undifferentiated “good” cells ^(16^. We selected a representative image (5120 x 5120 px) for each category from dataset A3, and obtained 1024 image patches (160 x 160 px) for each of the categories (Figure 4A). Then, the patches were processed by the trained VQ-VAE-2 model to extract the image features. We visualized the distribution of the extracted image features for each category by t-SNE dimension reduction (Figure 4B). In addition to that unsupervised analysis, we also evaluated the classification accuracy of the quality categories based on the extracted image features by a Support Vector Machine (SVM) model (Table 1). The image features were split 3:1 and used for training and test respectively. We obtain 89.3% test accuracy for the quality categories, after optimizing hyperparameters by 3-fold cross-validation for gamma and C with RBF kernel. Taken together, this analysis provides additional validation of our image features as accurate representations of iPSC quality, and an excellent agreement between VQ-VAE-2 model and expert curation.

**Figure 4A.**
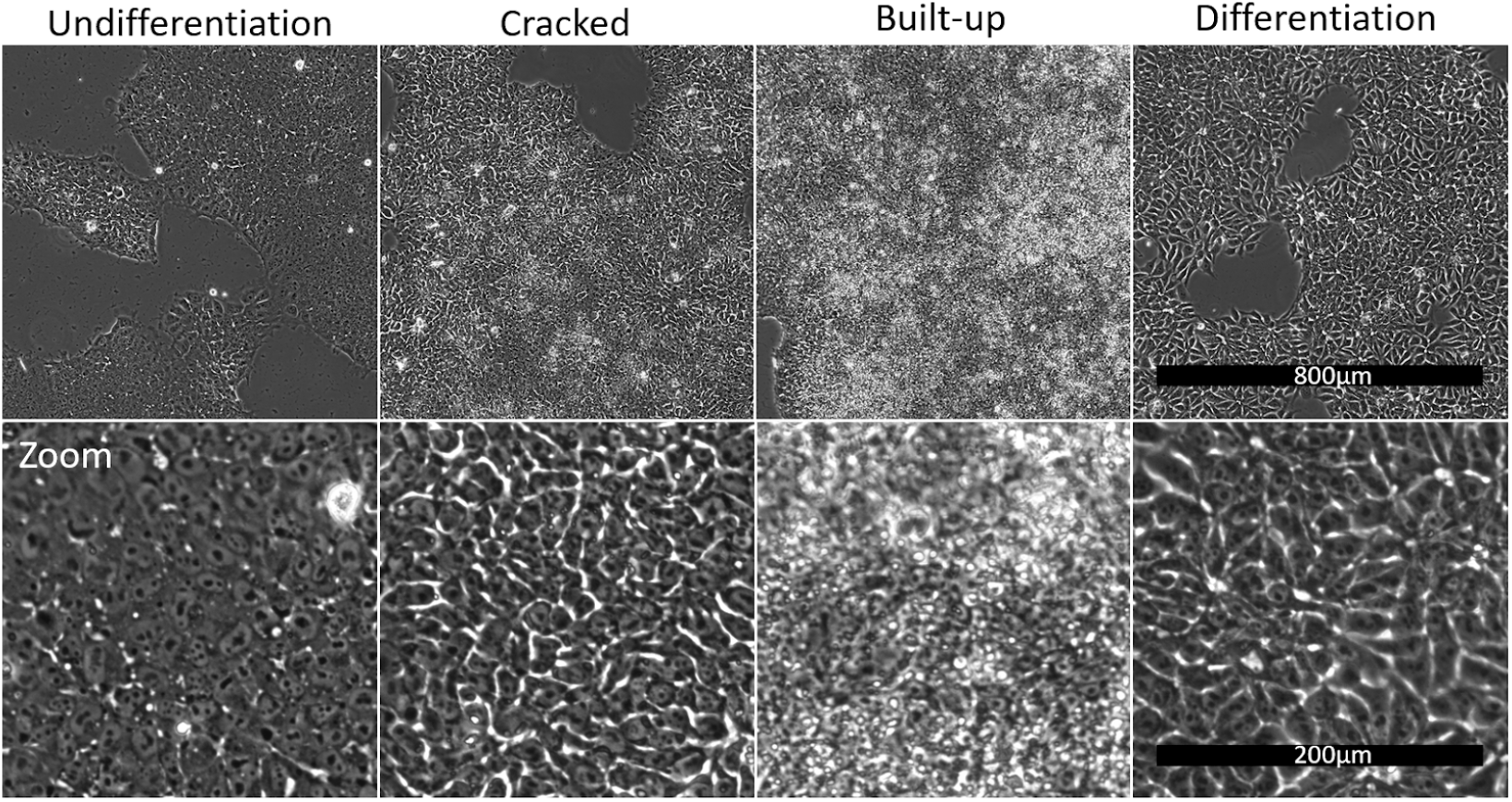
iPSC quality categories defined by the culture experts. Four morphological categories are visually defined by iPSC culture experts. *Undifferentiation:* Good iPSCs with high stemness and pluripotency. Cells have little cytoplasm and prominent nucleoli. *Cracked:* Upper-middle quality. Cells still have a morphology similar to undifferentiated cells, but some kind of differentiation-related activity appears as cracks. *Built-up:* Lower-middle quality. Cells are crowded, stacking on top of each other, and cell morphology cannot be observed at all. *Differentiation:* Low quality. Totally differentiated and is not useful for the subsequent differentiation process.

**Figure 4B.**
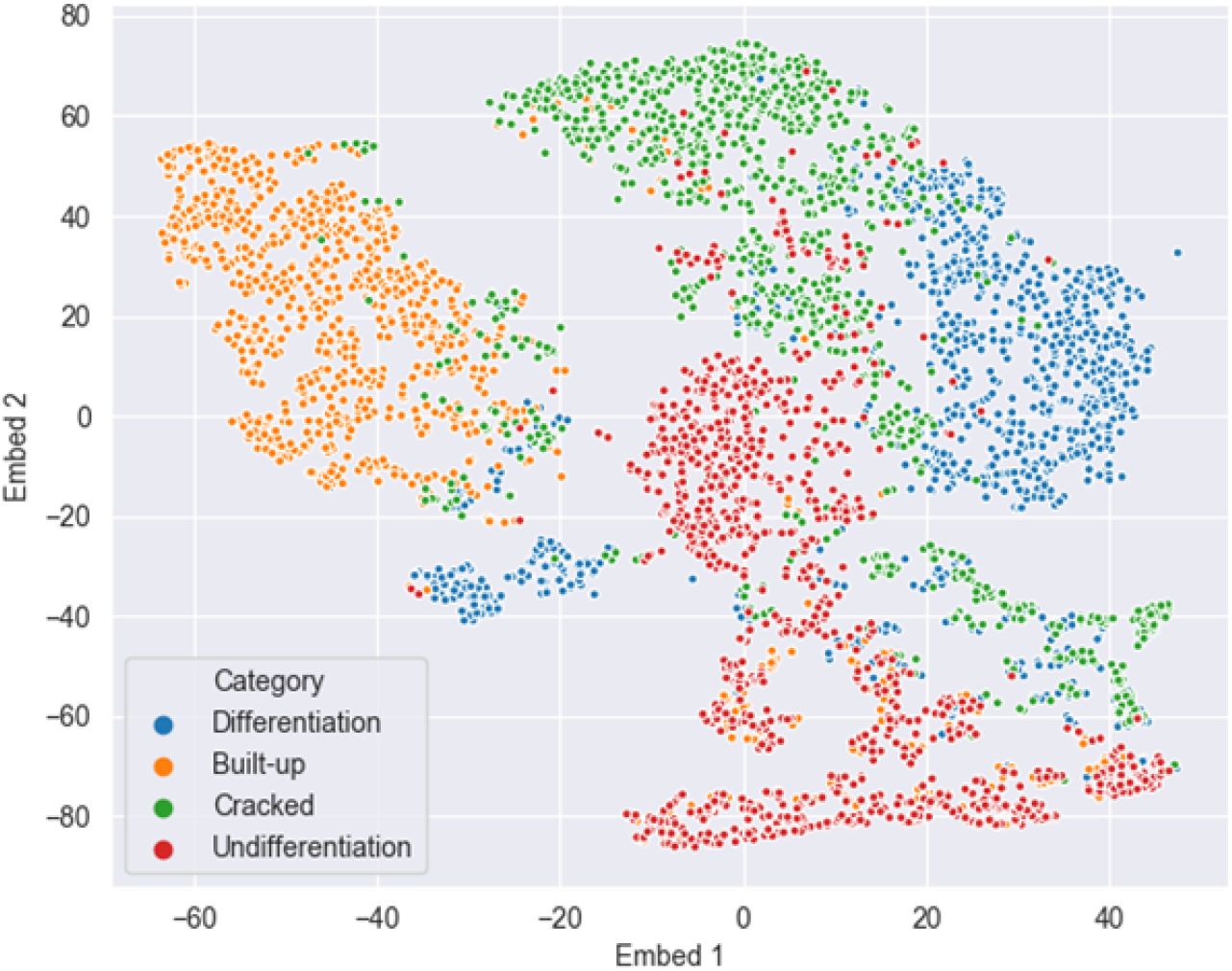
t-SNE plot of the image features extracted by the VQ-VAE-2 model trained with the A3 dataset. Each color represents the quality categories defined by the experts, which mostly exist in different regions in the feature space despite this being an unsupervised analysis which does not involve any training with expert annotations.

**Table 1.** Classification performance for the quality categories based on the image features.

|  |  | Number of patches | Confusion matrix |  |  |  |
| --- | --- | --- | --- | --- | --- | --- |
|  |  |  | Predicted category |  |  |  |
|  |  |  | Undifferentiation | Cracked | Built-up | Differentiation |
| True category | Undifferentiation | 255 | 234 (91.8%) | 13 (5.1%) | 4 (1.6%) | 4 (1.6%) |
|  | Cracked | 243 | 12 (4.9%) | 208 (85.6%) | 14 (5.8%) | 9 (3.7%) |
|  | Built-up | 249 | 4 (1.6%) | 11 (4.4%) | 233 (93.6%) | 1 (0.4%) |
|  | Differentiation | 277 | 4 (1.4%) | 29 (10.5%) | 4 (1.4%) | 240 (86.6%) |

### 2.5. Hierarchical clustering of gene expression levels and image features

To see the variance of the gene expression profiles across datasets, we performed hierarchical clustering analysis. We found that the gene expression profiles consisting of 218 genes have different characteristics between experiment A (one donor) and experiment B (14 donors) (Figure 5, left). We also performed hierarchical clustering of the image features across the datasets. This analysis revealed that image features of experiment A and experiment B don’t display clear clustering (Figure 5, right). This suggests that batch effects and their impact on gene expression profiles may pose a significant challenge to developing a single predictive model that would accurately fit all the datasets, and that multiple models may be needed.

**Figure 5.**
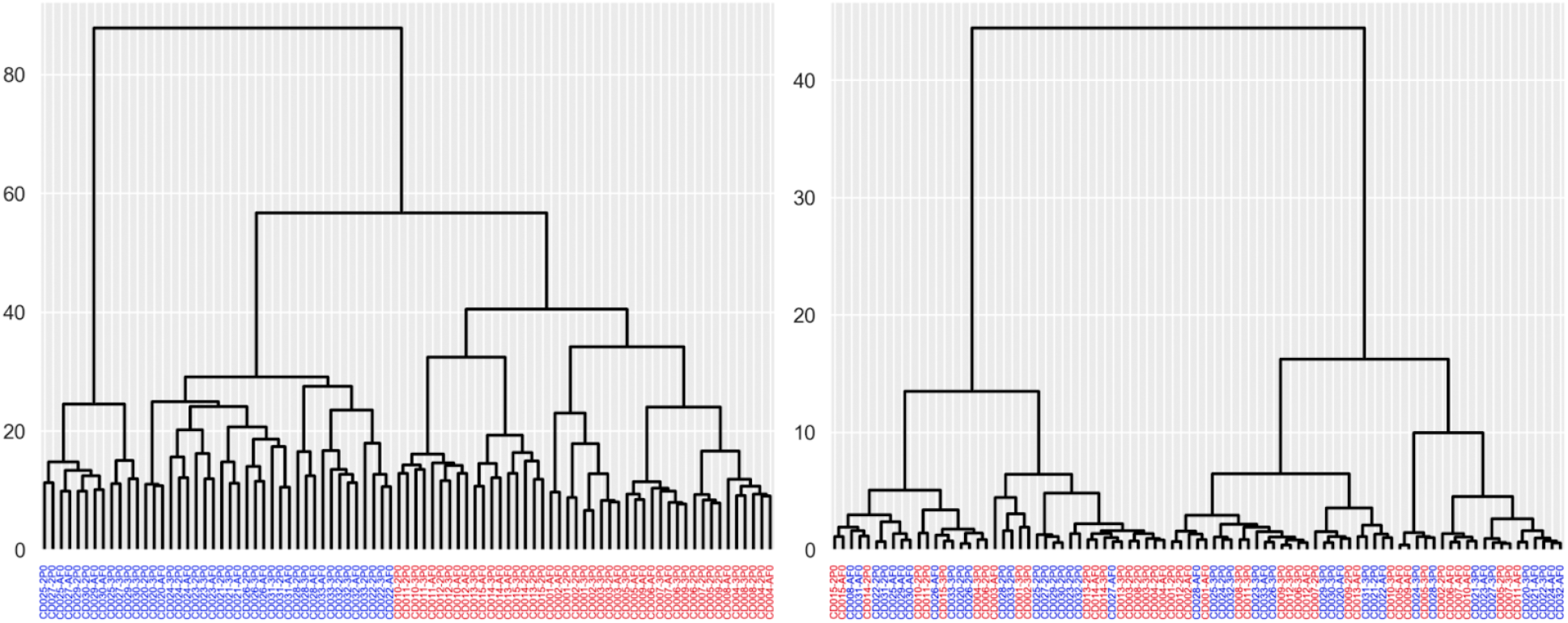
Hierarchical clustering results for the gene expression profiles (left) and the image features (right) across all 87 samples. Color represents the experiment. Blue: experiment A (clone #1-15). Red: experiment B (clone #16-29).

### 2.6. Validation with public benchmark dataset shows equivalent performance to an end-to-end approach

Before testing the VQ-VAE-based approach on the gene expression-from-brightfield-iPSC-images prediction task, we tested it on a distinct but related task – predicting gene expression using histopathologic images from a public spatial transcriptomics breast cancer dataset^(17^. This dataset contains 30,612 spots with gene expression profiles, in 68 breast tissue sections from 23 patients with breast cancer. The sections were scanned at 20x magnification, and 26,949 distinct mRNA expression levels were measured in spots with a diameter of 100μm arranged in a grid with a center-to-center distance of 200 μm. We compared the prediction performance of our model and the end-to-end CNN model DenseNet-121^(24^, which was previously used on this dataset, by focusing on the expression of the 250 genes with the highest mean expression for all the spots (validation methodology in Supplementary S1). The final prediction performance for each gene represents a median value of 23 correlation coefficients between measured gene expressions and predicted gene expressions for each patient’s single cells, obtained by leave-one-out cross-validation of 23 patients. As shown in Table 2, three of the top-five genes were consistent (p=3.8e-5, Fisher’s exact test) and our model performed comparably to the end-to-end CNN model ^(17^ (validation methodology in Supplementary S1). These results gave us confidence that our method can predict gene expression from images with accuracy comparable to the current state-of-the-art but without needing labels to learn image representations.

**Table 2.** Validation results on a published benchmark. Genes with top-5 prediction performances. Values in parentheses are the median of correlation coefficients between measured and predicted gene expressions obtained by leave-one-out cross-validation of 23 patients. Common genes are marked with an asterisk.

|  | DenseNet | Ours |
| --- | --- | --- |
| 1 | *HSP90AB1<br>(0.325) | *FASN<br>(0.390) |
| 2 | *FASN<br>(0.324) | DDX5<br>(0.372) |
| 3 | *GNAS<br>(0.319) | *HSP90AB1<br>(0.343) |
| 4 | ACTG1<br>(0.317) | TPT1<br>(0.314) |
| 5 | FN1<br>(0.315) | *GNAS<br>(0.309) |

### 2.7. Prediction of gene expression based on iPSC datasets

Having demonstrated the efficacy of the strategy on a benchmark task, we employed our strategy to predict gene expression values from extracted image features of iPSCs. We evaluated the prediction performance for all iPSC datasets using SVR models trained with either dataset A3 (one donor, timepoint 3) or B3 (14 donors, timepoint 3) (Table 3). In both cases, 3-fold cross-validation within the training dataset was performed to optimize SVR model hyperparameters for each gene as described in Methods. We then calculated the coefficient of determination (R^2^) and corresponding significance corrected by the Benjamini-Hochberg method (FDR), between the predicted gene expression and the measured gene expression for every dataset. Then, we picked the top-10 genes with the highest R^2^ value and that pass significance of α=0.05.

**Table 3.**
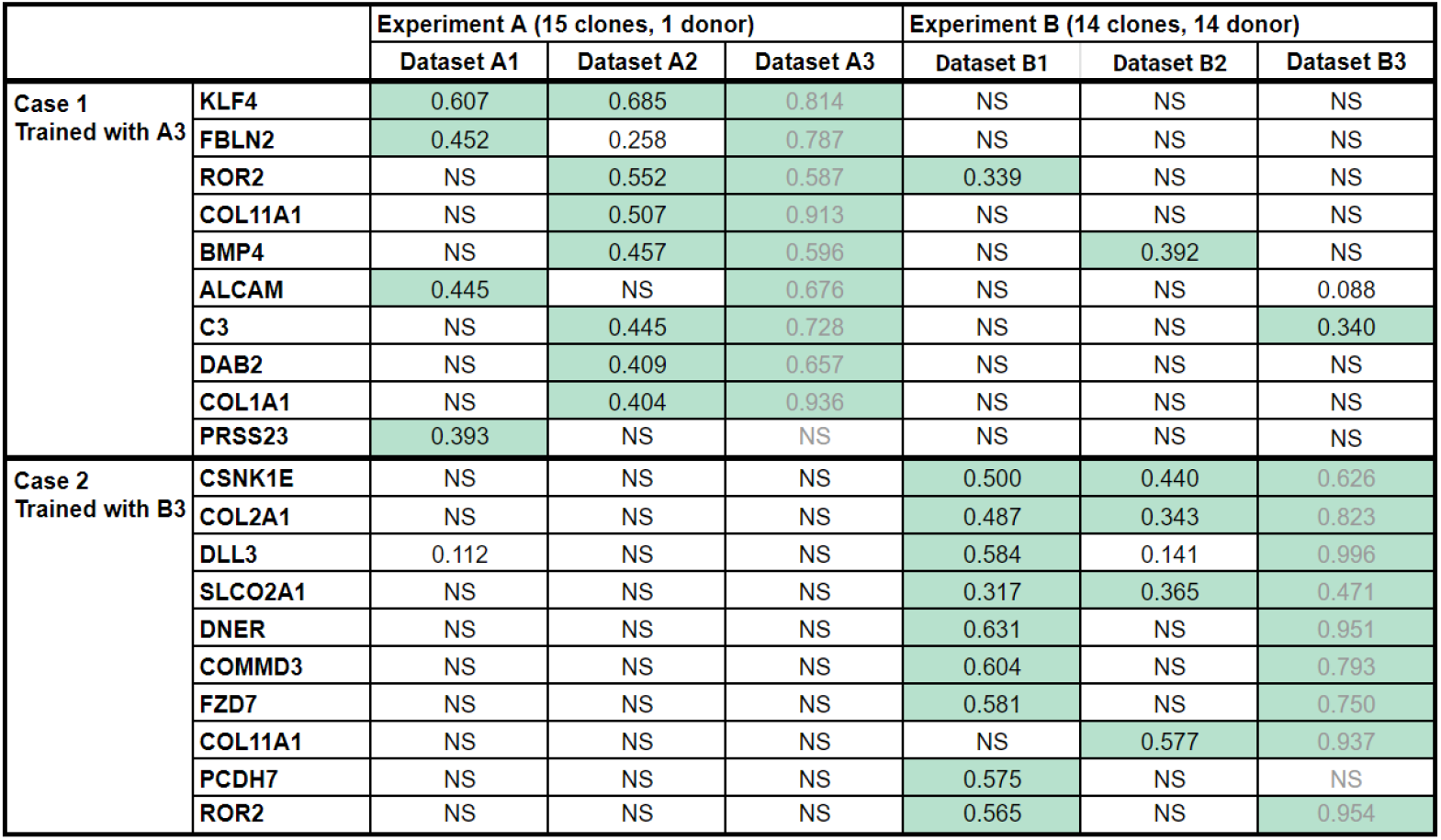
Prediction performance for the top 10 genes across experiments. Cells with R^2^ > 0.3 between predicted and measured gene expression levels are highlighted in green. NS: not significant with FDR correction (α=0.05). Results for the training set are in light gray.

When trained on the single donor dataset (A3, timepoint 3), the best-predicted gene was KLF4. For KLF4 (Figure 6A), the model obtained good predictions on the dataset A1 and A2 with R^2^=0.607 and 0.685 (and A3 used for training = 0.834) but did not perform well on the datasets of experiment B. This indicates that training does not generalize well across the experiments either due to technical variation (batch effects) or due to the differing donors. For the other genes, the model had R^2^ >0.3 for only one or two datasets other than the dataset used for model training.

**Figure 6.**
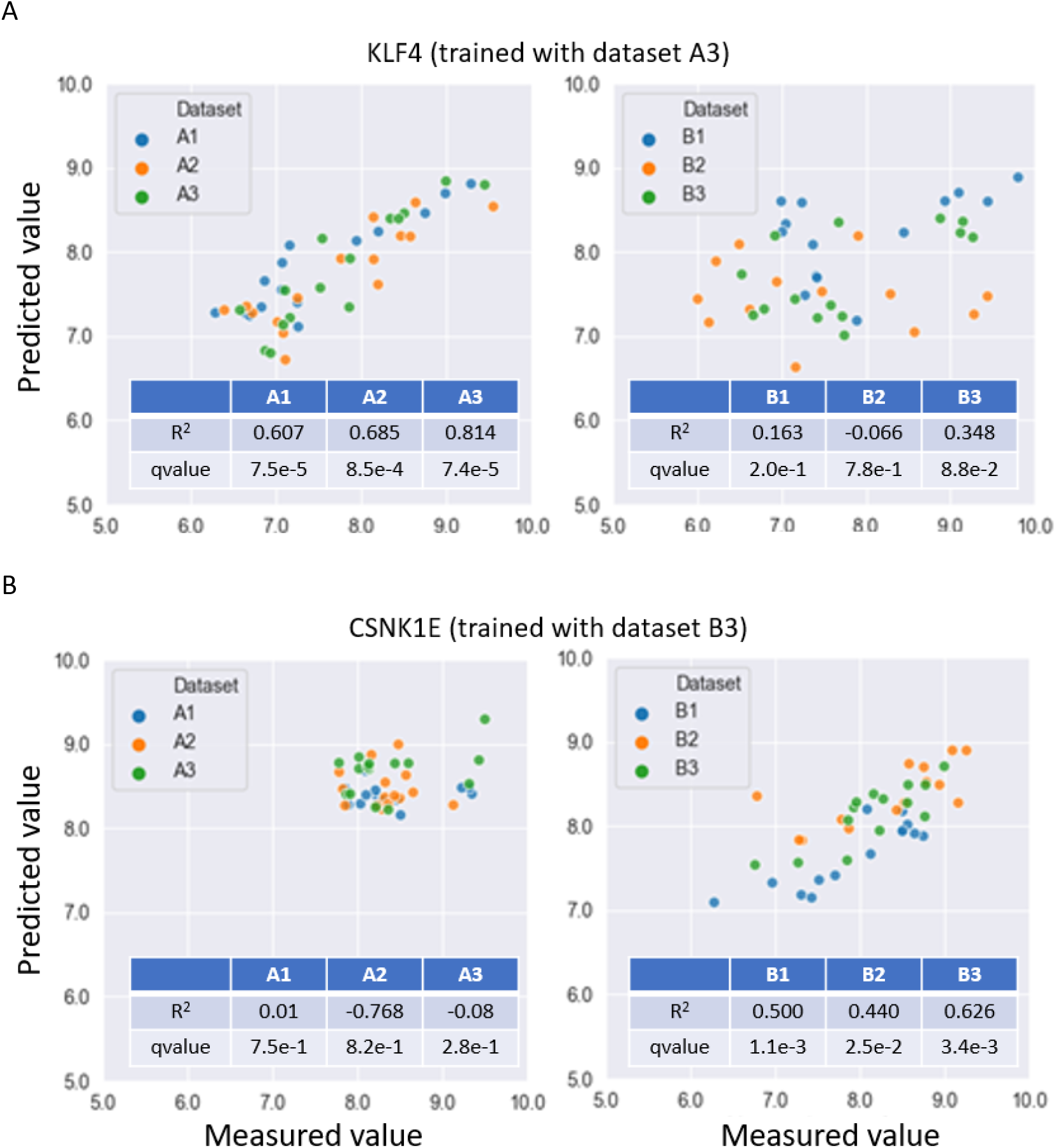
Gene prediction results for each dataset. Predicted (y-axis) versus measured (x-axis) expression levels (log2) for KLF4 (A) and CSNK1E (B) using the trained models as labeled.

We wondered whether training on a larger set of donors might improve predictions. We therefore trained on the 14-donor dataset (B3, timepoint 3), and found the best-predicted gene was CSNK1E. For CSNK1E (Figure 6B), predictions were good in datasets B1 and B2 with R2=0.500 and 0.440, again demonstrating good generalizability across timepoints within a batch. This model did not perform well on Experiment A datasets, however, which indicates that training on many donors does not improve generalization across the experimental batches. Nevertheless, training with 14 donors (as compared to single-donor Experiment A) did show an advantage: it yielded stronger ability to predict genes within the experimental batch, especially for dataset B1.

To investigate how well specific morphological characteristics correspond to best-predicted gene expression levels, we collected patch images from the datasets according to the gene expression level of the best-predicted gene for each patch. We observed that the cells show differentiation-like morphology when the predicted expression level of KLF4 is low, and change into the undifferentiation-like morphology as the predicted KLF4 expression level increases, for images within Experiment A (Figure 6C). For images within Experiment B, we also confirmed that as the predicted CSNK1E expression increases, the cells initially change their appearance from built-up-like morphology to undifferentiated-like morphology, until they become larger and the cell boundaries become unclear (Figure 6C). Taken together, although the noted issues with achieving consistent predictability points to batch-specific technical variation or donor-specific variation as main factors that impact prediction performance, morphological analysis suggests that changes in image features correlate with changes in predicted expression levels for best-predicted genes.

**Figure 6C.**
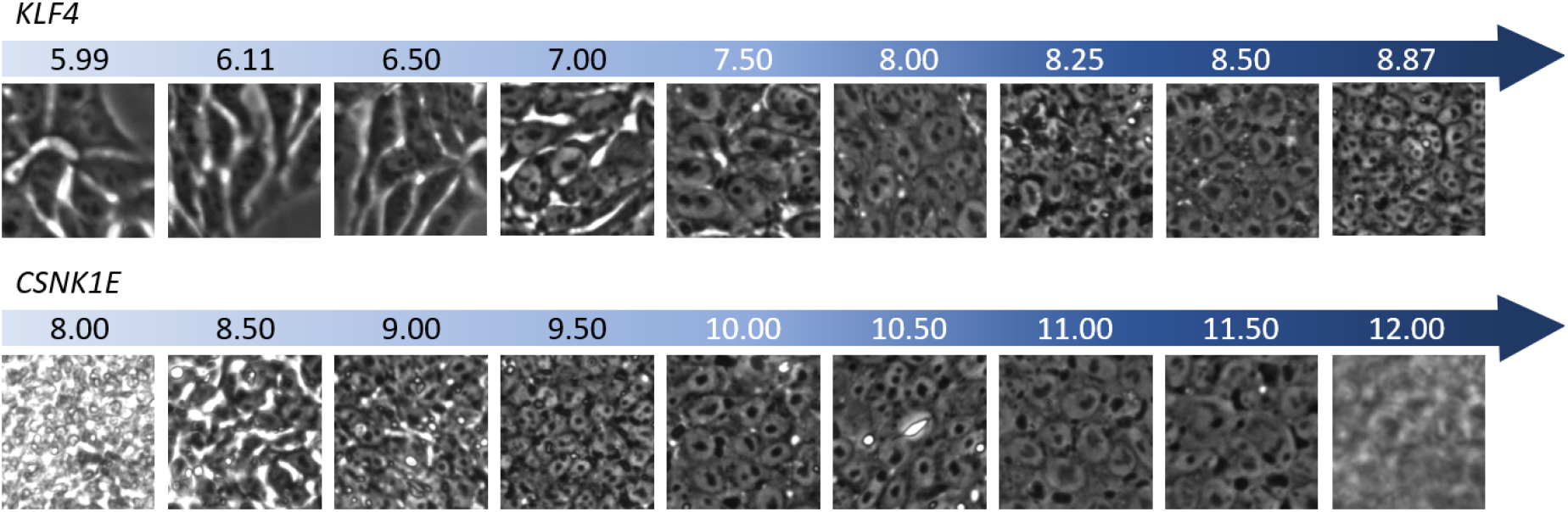
Cell morphology and predicted gene expression levels for KLF4 and CSNK1E. The numbers represent the predicted gene expression levels corresponding to each image. The patches are collected from the datasets of experiments A and B for KLF4 and CSNK1E, respectively.

## 3. Discussion

In this study, we developed a strategy aimed at improving performance of the iPSC purification process by using cell morphology captured by phase-contrast microscopy to predict gene expression levels relating to stemness and pluripotency. To achieve this, we introduced a new method to predict gene expression levels based on label-free images of cells that employs a classifier trained on image features extracted using an unsupervised deep learning model, VQ-VAE-2. Furthermore, unlike all other currently available strategies that rely on single cell RNA-seq, a method that is destructive to the entire sample under analysis, we used bulk-level gene expression profiles that can be obtained for partially sampled cells so that cells can continue to be cultured after the gene expression measurement.

We benchmarked the performance of our method using a completely different setup – a public dataset of tissue slide images and spatial gene expression data – and demonstrated that the method performs as well as the state-of-the-art while requiring much less labeled data. We also investigated the relationship between the image features captured by our method and the iPSC morphological quality categories experimentally defined by the iPSC culture experts. The results showed that there is a correspondence between the morphological quality categories and the image features extracted from unstained phase-contrast images, allowing morphological quality categories to be predicted from the image features with 89.3% accuracy. Although the relationship between the morphological categories assigned by human experts and gene expression levels are unclear, our results suggest that the method we developed could assist in linking image features with both gene expression and the quality categories. This would further facilitate visual inspection and iPSC purification, and make the process more reliable.

When we employed our approach to predict gene expression from image features, we identified two top-predicted genes, KLF4 and CSNK1E, from two different experiments. For KLF4, we observed that as the predicted expression increases, the appearance of cells changes from differentiated-like to undifferentiated-like. This observation agrees with the known biology of KLF4, as it has previously been established that KLF4 is a key gene for iPSC reprogramming and de-differentiation process in somatic cells, and that this process can be paused by manipulating the KLF4 expression ^(25–26^. For CSNK1E, we observed that as the predicted expression increases, the cellular morphology changes from built-up-like, to flat, undifferentiated-like, to having an unclear cell boundary, indicating that the cells were progressively dissociating. This observation appears consistent with CSNK1E biology as this protein accelerates cell migration and cell dissociation. ^(27–28^ Therefore, the gene expression predictions from image features for top genes are in good agreement with their established biological functions.

Our results also highlight that we obtained two significantly different predictions in two different experiments, Experiment A that included 15 clones from the same patient, and Experiment B that included samples from 14 different patients. This suggests that batch effects, i.e. non-biological factors in an experiment that are observable in the data and known to pose a major challenge in high-dimensional biology ^(29^, play a significant role in our work as well. We observed that the predictive model works well only when the training and test dataset are from the same batch (i.e. same experiment); training a model on many donors (Experiment B) did not dramatically improve predictions vs training on a single donor. We also observed that the list of genes with the best R^2^ values is different across the batches. This suggests that not only are the models non-generalizable across the batches but also the relationships between morphology and gene expression appear to be different across the two batches. This discrepancy is consistent with the observation from our hierarchical clustering analysis, where we noted that strong batch effects are observed in gene expression data but not in the imaging data.

In our experience, batch effects are predominantly due to the stochastic nature of the iPSC reprogramming process itself^13,4^. In fact, the iPSCs for the two experiments (A and B) were reprogrammed separately, so it is very likely that the batch effects are indeed driven by the reprogramming process. Given this source of batch effects, our overall approach can be used to make predictions of expression levels only within a reprogramming batch. Despite this current caveat, our methodology can be usefully implemented by collecting data and training a model early in the iPSC reprogramming process and applying it to future timepoints of the batch, thus eliminating repeated gene expression profiling for the many subsequent rounds of cloning. Furthermore, batch-effect correction techniques applied to such data, together with a more diverse set of training data, may further improve the ability to generalize the models, reduce the need for gene expression profiling, and enable automated reprogramming processes based on live cell imaging.

In conclusion, we explored the relationship between iPSC morphology and gene expression within iPSC multi-clone datasets collected during the simulated iPSC purification process. We used phase-contrast images as a morphology readout and bulk RNA-seq as gene expression data, and trained the models to use image features to predict gene expression. Although batch effects, driven by the iPSC reprogramming process, limit our ability to generalize the model across reprogramming batches, the proposed approach may allow stem cell facilities to automatically track the iPSC purification process using phase-contrast imaging alone.

## 4. Methods

### 4.1. Data acquisition

In this study, we used six datasets composed of 29 iPSCs clones derived from peripheral blood mononuclear cells (PBMCs) of 15 donors. The culture of iPSCs was carried out in two phases. We first cultured 15 iPSC clones (Experiment A) all derived from a single donor, and then cultured 14 iPSCs clones derived from 14 donors (Experiment B), one clone per donor, using the same iPSC culture protocol ^(4^. To simulate the iPSC purification process, all iPSCs were cultured for three passages with different types of vessels, 6-well plate, T75 flask, and T150 flasks for each passage respectively (Supplemental Fig.1). In every passage, the cells were passaged on Day 4, and phase-contrast imaging and gene expression measurements were performed before passaging. As a result of these culture experiments, we obtained six datasets composed of 29 clones and three time points (87 samples, Supplemental Table 1).

### 4.2. Phase-contrast imaging

The images were obtained by a phase-contrast microscope (Nikon Ti2-E) with a 10x objective lens(Nikon Plan Fluor 10x/0.30 DL and a 25M pixel camera (25CXP1, CIS). We captured single-focal-plane grayscale images of cells for each culture vessel. We captured 60, 100, and 100 images for Timepoint 1, Timepoint 2, and Timepoint 3 respectively. The resolution and the dimensions of each image are 0.45 um/px and 5120×5120px respectively.

### 4.3. Gene expression normalization method

To enable comparison of the expression levels between samples, measured expression profiles were normalized by the total number of reads for each sample. Then, we took a logarithm of base 2 after adding a pseudo count of 1 to prevent zeros.

### 4.4. VQ-VAE-2 model training

We trained a VQ-VAE-2 model as it is, with K of 64 and D of 64 on the images of a single dataset (A3) using Adam optimizer ^(23^ with a learning rate of 1e-4 to minimize MSE between the input patch and the restored patch. All the images included in A3 were split into small patches of 160×160 px and used in the model training. The training iterated for 50 epochs, with batch size 32. We used the PyTorch machine learning framework for implementation.

### 4.5. Extraction of image feature vectors

We extracted the latent feature vector of a sample by the following procedure. First, we split all the images of a sample into small patches. Image size is 5,120×5,120px and we adopted a patch size of 160×160 px, so the number of patches per image is 1,024. Each patch contains roughly 30 cells. Second, we extracted vector-quantized feature maps by applying the trained VQ-VAE-2 model to all the patches. Here we have two feature maps with different resolutions (20px and 40px each) per patch. Third, we counted the appearance frequency of each embedding vector from the feature maps for all the patches. We totaled these and normalized the total frequency of embedding vectors by each resolution after adding a pseudo count of 1 to prevent zeros from being used in the next step, and then computed the logarithm of base 2, to get the sample’s profile.

### 4.6. Gene expression prediction model training

We applied SVR to predict the gene expression of each sample from sample-level image features using the scikit-learn machine learning library in Python. Training of SVR models and optimization of the model hyperparameters were done for each gene, in other words, we constructed as many SVR models as target genes (n=218). For SVR models, the linear kernel function was adopted, and hyperparameters (C, epsilon) were searched from the following lists by a grid-search algorithm with 3-fold cross-validation in the training dataset:

- C: [1,2,3,…,100]
- epsilon: [0, 0.05, 0.10,…,0.50]

### 4.7. Hierarchical clustering

Hierarchical clustering was performed using linkage function of Scipy library. The Ward method was used to calculate distances. For images, clustering was performed on the image features of each sample. For gene expression, clustering was performed for the expression profiles of the extracted 218 genes after normalization.

### 4.8. Morphology feature association study

Gene prediction was performed for 10 representative images selected from each sample. The VQ-VAE-2 model trained with the images of a single dataset(A3) was applied to all patches of the selected images to obtain image features for each patch. Then the SVR models to predict genes were applied to the image features. Each SVR model was applied only to the image features calculated from the dataset which the model was trained with. The KLF4 prediction model was applied to the image features obtained from the dataset of Experiment A. The CSNK1E prediction model was applied to the image features obtained from the dataset of Experiment B. To obtain gene expression levels patch-wisely, the feature averaging step was omitted.

### 4.9. Data and code availability

We used the VQ-VAE-2 implementation from https://github.com/rosinality/vq-vae-2-pytorch and did not make any modifications. All the relevant parameters for model training and feature extraction are documented above.

## 5. Acknowledgments

AEC acknowledges support from the National Institutes of Health (NIH R35 GM122547 to AEC)

## Supplementary Materials

### 1. External validation process

As mentioned in section 3.3, we tried to validate our method on an external spatial transcriptomics dataset. We concluded that our method, using the VQ-VAE-based image feature extractor, performed at least equivalent to the end-to-end CNN model in gene expression prediction from image. In this supplemental section, we described the details of the validation process.

We first implemented an end-to-end model to predict expressions of 250 genes from small histopathologic image patch by following the description in the paper ^(17^. DenseNet-121 ^(24^ model (pretrained by ImageNet dataset) followed by a dense layer with 250 units was trained as a retest model using 224×224 image patches and corresponding gene expressions of the spots. We modified our model to solve the same task. We trained a VQ-VAE-2 model in advance and its encoder was used as an image feature extractor. Instead of SVR, a MLP model was trained as a gene expression predictor using 224×224 image patches and corresponding gene expressions of the spots. VQ-VAE-2 hyper parameters were not changed (K=64, D=64). The MLP model has 3 layers: (1) input layer with 128 units, (2) hidden layer with 1,024 units (same as the output dimensions of DenseNet-121), (3) output layer with 250 units (See Fig. S1). Other training configurations below were applied both the retest model and our model:

- Loss: MSE
- Optimizer: stochastic gradient descent with learning rate of 1e-6 and momentum of 0.9
- Epochs: 50
- Batch size: 32
- Data augmentation: randomly rotating the image by 0, 90, 180 or 270° and taking the mirror image 50% of the time

**Fig.S1.**
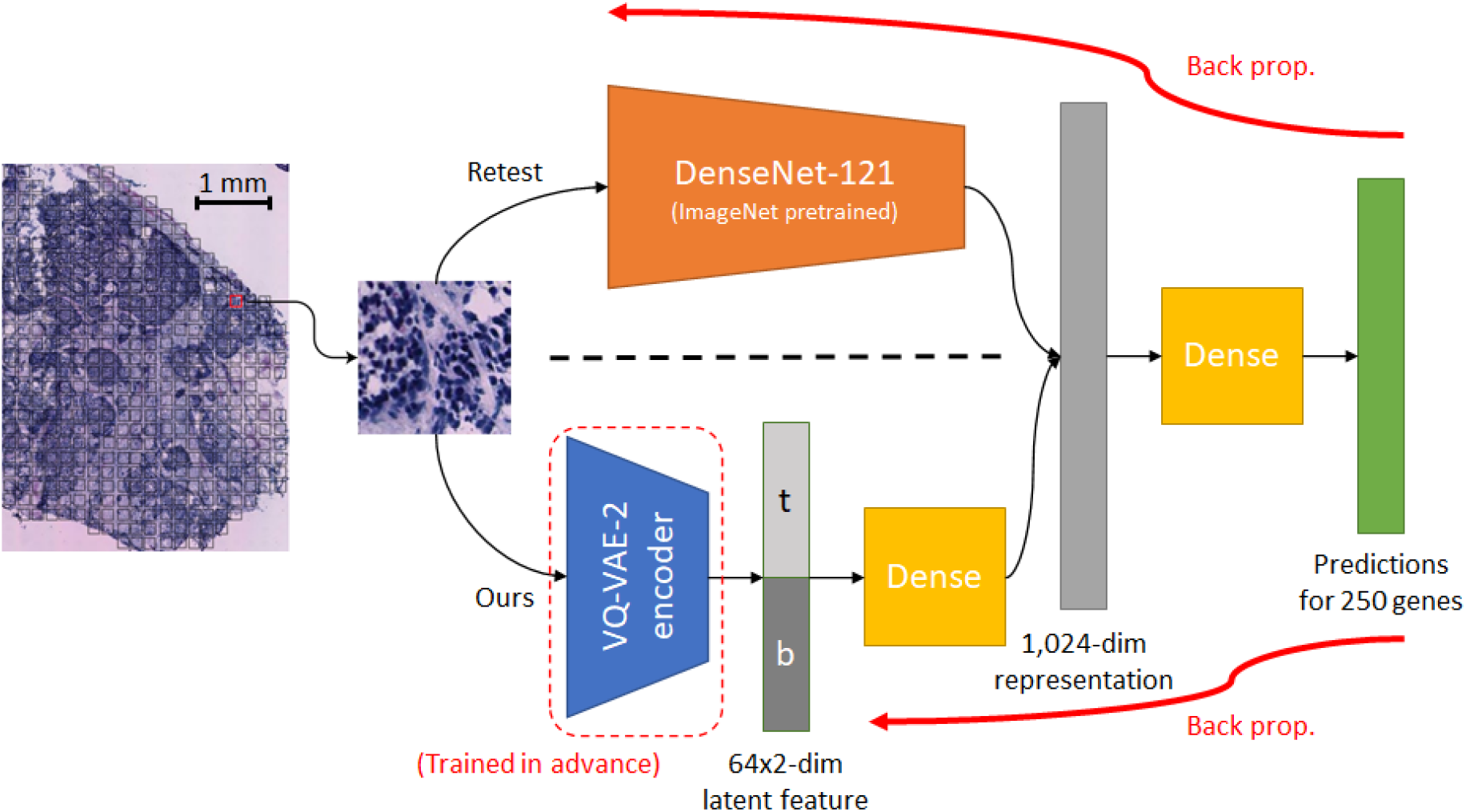
Model training for external validation. The retest model (DenseNet-121) is trained in the end-to-end manner to minimize MSE for gene expression prediction. On the other hand, our model uses freezed VQ-VAE-2 encoder as a feature extractor, and the simple MLP model is trained to minimize MSE.

Then we compared the results of the rest model and our method. We evaluated models by one-leave-out cross validation of 23 patients. As mentioned in section 3.3, the prediction performance for each gene is a median value of 23 correlation coefficients obtained by the cross validation. As shown in Table S1(a), although the ranges of top-5 prediction performances were smaller than the paper (left column), 3 of top-5 gene names were matched (p=3.8e-5) in both retest model and our model. Table S1(b) shows the prediction performance rankings of top-5 genes in the paper and 4 of top-6 gene names were matched except for XBP1.

**Table S1.**
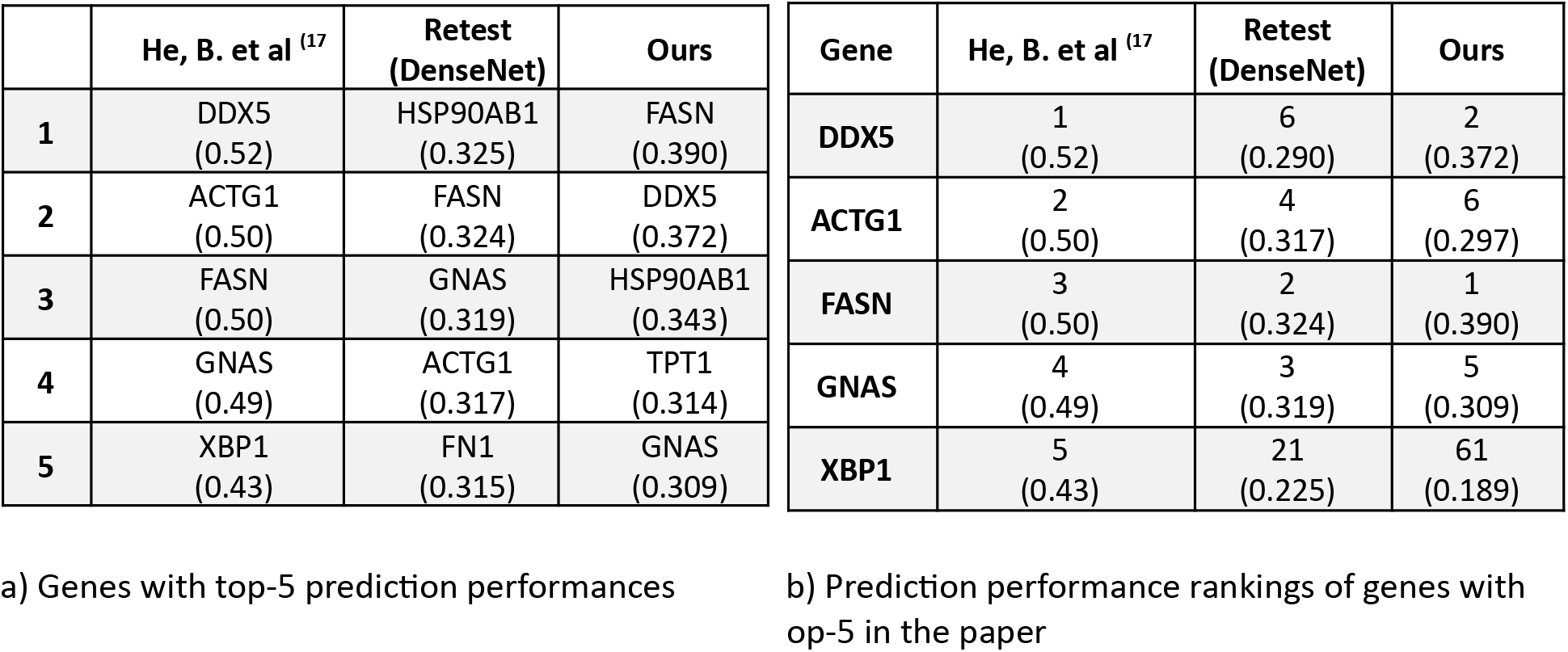
Results of ST-Net retest and our method validation. Values in () are median of correlation coefficients between measured and predicted gene expressions obtained by one-leave-out cross validation of 23 patients.

We could not reproduce the result of the previous study completely, but the trends are consistent. Therefore, we considered that we do not need to change our conclusion that our model works as well as the end-to-end model. On the other hand, the inadequate reproduction might be caused by newly implementing the retest program instead of using their official sources. We have noticed (but not confirmed so far) that potentially there were several mismatches between the original and ours such as 250 genes for prediction targets (they extracted 250 genes with highest mean expressions as prediction targets, but the entire list of them was not found.) and data loading (in their source, missing table data were referred in order to load spatial gene expressions and corresponding spot IDs.).

### 2. Restoration result by VQ-VAE2 model

**Fig.S2.**
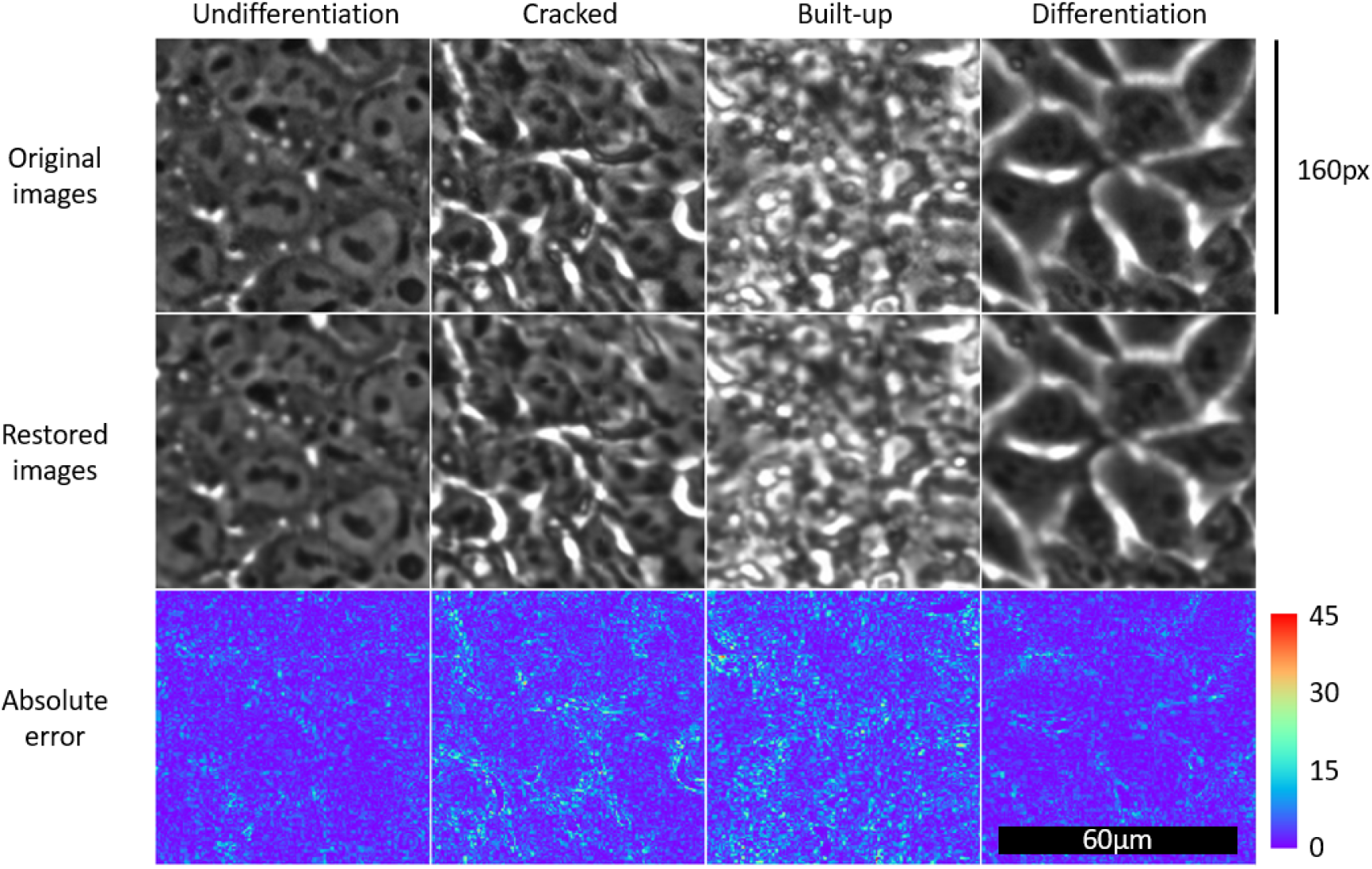
Comparison between original images and restored images for each iPSC quality category. Color bar represents absolute error level in 8bit between the original and the restoration. A little difference between the images can be found mainly in high frequencies so that the restored images tend to have more smooth appearance than original images.

